# Coacervate Droplets as a Liquefier for the Solid-to-Liquid Transition of RNA Aggregates

**DOI:** 10.64898/2026.01.18.700211

**Authors:** Wei Guo, Rong Luo, Yi Shen, Xiangze Zeng, Zhou Liu, Ho Cheung Shum

## Abstract

Biomolecular condensates are functionally gated by their material properties. While RNA is a primary structural scaffold, its sequence-specific interactions can aberrantly drive condensates into dysfunctional solid aggregates. Yet, actively reversing this solidification to restore function remains a fundamental challenge, impeding progress in synthetic biology and therapeutics interventions. Here, we establish that complex coacervates can be engineered as liquefiers to actively remodel solid RNA-peptide aggregates into functional liquid droplets. Integrating systematic experiments with all-atom simulations, we decode a multiscale mechanism: coacervate infiltration at the micrometer scale mediates molecular buffering at the nanometer scale, which fluidizes the crosslinked network to drive macroscopic liquefaction. This capability is governed by a design rule, where coacervates formed by small-molecular anions exhibit optimal efficacy. We deploy this principle to functionally rescue silenced RNA within a model of pathologically solidified ribonucleoprotein assemblies. Our work provides a general framework for the active, compositional control of biomolecular phase behavior, with direct implications for managing pathological aggregation and engineering functional condensates in synthetic and living systems.

## Introduction

Biomolecular condensates formed through liquid-liquid phase separation (LLPS) serve as dynamic, membraneless compartments that spatially organize biochemistry within cells^1^. The material properties of these condensates are tunable, spanning a programmable spectrum from liquid-like droplets^2^ to solid-like aggregates^3, 4^. The progression of liquid-to-solid transition of biomolecular condensates, termed ageing, drives the formation of dysfunctional aggregates implicated in neurodegenerative disease^5^. Actively remodeling biomolecular aggregates to restore function, however, represents a fundamental challenge and a critical frontier in the material control of condensates^6–8^.

Both proteins and RNAs contribute to the aberrant aggregation of condensates^9–11^. For proteins including hnRNPA1^12^, FUS^13^, and Tau^14, 15^, ageing accompanies cross β-sheet formation that drives a thermodynamically stable, often irreversible conversion into gel– or amyloid-like aggregates. This irreversibility confines strategies to the prophylactic manipulation of the ageing pathway prior to aggregation, using agents like ATP hydrotrope^16^, molecular chaperones^17^, or peptide inhibitors^18^. In parallel, RNAs with GC-rich repeats^19, 20^ and noncanonical structures^21^ are intrinsically prone to form solid-like aggregates, critically implicated in Huntington’s and amyotrophic lateral sclerosis^19, 20^. This aggregation can be driven by homotypic RNA-RNA interactions^22,23^ or by heterotypic interactions with RNA-binding proteins (RBPs) such as ribonucleoproteins (RNP)^24^. Crucially, a key aggregation type involves strong multivalent interactions that kinetically trap assemblies into a solid-like state^25–27^. These kinetically arrested assemblies do not reside in a deep free-energy minimum, implying a latent thermodynamic accessibility to a functional liquid state, even after aggregation. This opens a distinct, unexploited paradigm for condensate engineering, establishing the reversal of aberrant aggregation as a tangible objective for material control.

Recent findings demonstrate that the internal milieu of biomolecular condensates can mimic organic solvents, exhibiting low dielectric permittivity^28^ and a propensity to destabilize nucleic acid secondary structures^29^, despite their primarily aqueous composition. Particularly, the presence of condensates could drastically affect protein and RNA aggregation process^7^. For example, protein aggregation can be accelerated^30^ or suppressed within condensates^31^, or specifically enhanced at the interface^32, 33^. Most recent discoveries highlight that RNA aggregation is enhanced within multicomponent condensates, whereas introducing RBPs can disassemble these aggregates through competitive heterotypic interactions^34^. This duality demonstrates that the internal interaction network of a condensate can directly regulate the aberrant crosslinking that led to aggregate formation. These insights converge on a pivotal question: can the intrinsic environment of a condensate be harnessed as a solvent to liquefy solid-like RNA assemblies and restore their functions?

Here, we report that specific biomolecular condensates function as an effective liquefier for the solid-to-liquid transition of RNA-peptide aggregates. As a model for disease-relevant solids, we employed GC-rich RNAs prone to form inter– or intramolecular hybridization, to complex with cationic peptides. The resulting aggregates are amorphous solid-like condensates arrested by strong RNA-peptide interaction networks^21^. We demonstrate that liquid-like condensates, particularly those formed by small anionic molecules (such as ATP, UTP or ADP), can serve as liquefiers to dissolve these aggregates. Integrating multiscale experiments and simulations, we establish that condensate-mediated infiltration of small anionic molecules buffers the kinetically trapped interaction network within the aggregate, driving the solid-to-liquid transition. By harnessing this principle, we demonstrate that condensate-mediated liquefaction can reactivate encapsulated RNA function within a synthetic model of pathologically solidified RNP assemblies. Our work uncovers a novel regulatory function of biomolecular condensates: the active buffering of RNA phase transitions through solvent-like intervention. This provides a transformative framework for the rational engineering of biomolecular condensates as liquefaction medium, unlocking previously inaccessible therapeutic strategies for diseases driven by aberrant RNA aggregation.

## Results

### Coacervate droplets trigger the solid-to-liquid phase transition of RNA-peptide aggregates

Our investigation was prompted by a striking macroscopic phenomenon: the introduction of the mixture containing pre-formed liquid-like coacervate droplets to solid-like RNA aggregates induced a complete phase transition (Fig. 1A and Supplementary Fig. 1). Upon contact and infiltration by coacervate droplets (Supplementary Fig. 2), the initially irregular aggregates undergo a progressive liquefaction, transitioning into spherical, liquid-like condensates (Fig. 1B, C). These resultant droplets exhibited classic liquid characteristics, including wetting on a glass surface (Fig. 1B), suggesting a solid-to-liquid phase transition. Solid-like RNA aggregates were formed via electrostatic complexation of poly-L-lysine (PLL) with rPU22, a G-quadruplex-forming RNA oligonucleotide with sequence context relevant to GC-rich repeat expansion disorders, as previously reported^21^ (Supplementary Fig. 3), while the liquid coacervates were formed from ATP (adenosine triphosphate) and PLL (Fig. 1D, E). Crucially, the RNA solid aggregates are stable and not dissolved by increased ionic strength, confirming that simple charge screening cannot replicate the liquefying action of the coacervates (Supplementary Fig. 4). Fluorescence imaging confirmed that the transformed condensates incorporated the fluorescently labelled rPU22 RNA, indicating that the RNA from the original aggregates remained a core component of the new liquid phase.

**Figure 1.**
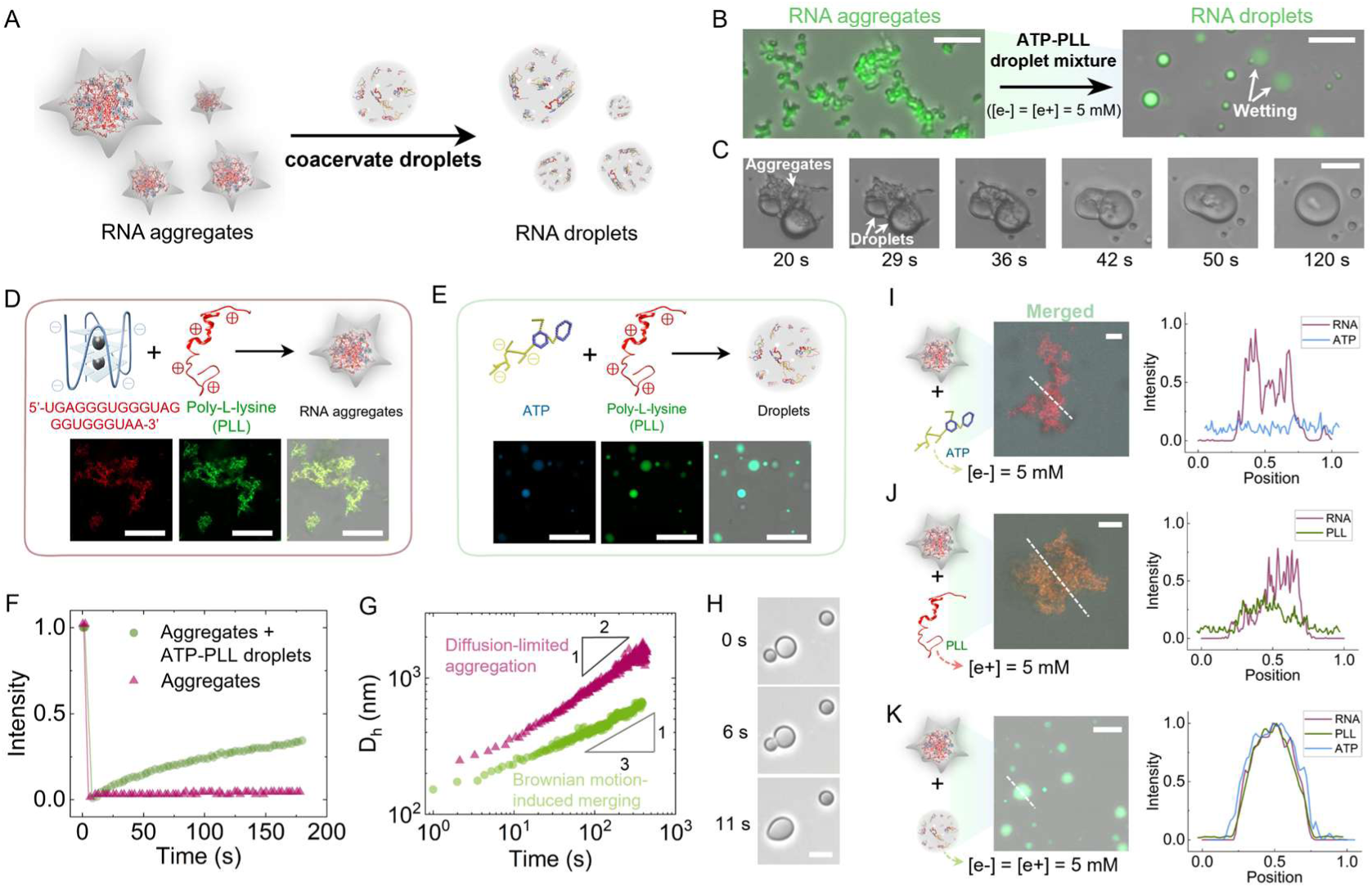
| ATP-PLL coacervate droplets trigger a solid-to-liquid phase transition of RNA-peptide aggregates. (**A**) Schematic illustration of the coacervate-mediated phase transition of solid RNA aggregates into liquid droplets. (**B**) Fluorescence micrographs show the macroscopic transition, where FAM-labelled RNA within aggregates also exhibits its integration into the new liquid droplet phase. Scale bar: 50 µm. (**C**) The solid-to-liquid transition of RNA aggregates upon contact (t = 0 s) with coacervate droplets. Scale bar: 20 µm. (**D**) Solid-like RNA-PLL (Poly-L-lysine) aggregates formed through G-quadruplex-mediated complexation. Cy5-labeled RNA and FITC-labeled PLL were used for fluorescence imaging. Scale bar: 20 µm. (**E**) ATP and PLL form liquid-like coacervate droplets. ATTO-390-labeled ATP and FITC-labeled PLL were used for fluorescence imaging. Scale bar: 20 µm. (**F**) Fluorescence recovery after photobleaching (FRAP) analysis of the initial RNA-PLL aggregates and the transformed droplets. (**G**) Hydrodynamic diameter growth kinetics of the initial RNA-PLL aggregates and the transformed droplets. (**H**) Time-lapse micrographs showing the coalescence of two transformed droplets. Scale bar: 5 µm. (**I**) Free ATP (ATTO-390 labelled) is added to RNA-PLL aggregates (Cy5-RNA added). No uptake of ATP or phase transition occurs. (**J**) Free PLL (FITC labelled) is added to RNA-PLL aggregates (Cy5-RNA added). PLL incorporates but does not induce fluidization. (**K**) Pre-formed ATP-PLL coacervates (ATP: ATTO-390 labelled, PLL: FITC labelled) are added to RNA-PLL aggregates (Cy5-RNA added), leading to component mixing and the formation of homogeneous liquid droplets. Scale bars in (**H**)-(**J**): 10 µm.

We then quantitatively characterized the material properties of the initial and transformed condensates. Fluorescence recovery after photobleaching (FRAP) analysis on fluorescently labeled RNA within aggregates revealed that the original rPU22-PLL aggregates were solid-like, exhibiting negligible fluorescence recovery. In stark contrast, the droplets formed after the addition of ATP-PLL coacervates showed rapid and significant recovery, confirming their liquid character and high internal molecular mobility (Fig. 1F and Fig S5). Further evidence was provided by monitoring condensate growth kinetics via dynamic light scattering (DLS). The hydrodynamic diameter (*D*_h_) of the rPU22-PLL aggregates grew with a time exponent α ≈ 1⁄2, consistent with a diffusion-limited aggregation mechanism typical of solids^35^. Conversely, the transformed droplets grew with α ≈ 1⁄3, a scaling law characteristic of liquid-like coacervates that evolve via Brownian motion-induced coalescence (Fig. 1G). Finally, the rapid coalescence of these new droplets into larger structures (Fig. 1H) provided direct visual confirmation of their liquid nature. Collectively, these data unequivocally demonstrate that ATP-PLL coacervates enable the liquefaction of solid RNA-PLL aggregates into dynamic, liquid-like droplets.

We next asked whether this transition was a specific property of the pre-formed ATP-PLL coacervate droplet, or a nonspecific effect of its individual components. Control experiments revealed that adding free ATP or PLL solution at concentrations equivalent to those in the ATP-PLL coacervate mixture failed to induce any phase transition of the solid aggregates persisted (Fig. 1I, J). In contrast, the ATP-PLL coacervate mixture consistently triggered the transformation (Fig. 1K). Fluorescence imaging provided mechanistic insight: the transformed droplets incorporated both fluorescently labelled ATP and PLL from the coacervates (Fig. 1K), while no uptake of fluorescently labelled ATP was observed when adding ATP alone into RNA aggregates (Fig. 1I). Interestingly, adding free PLL to the aggregates increased the intra-aggregate fluorescence signal but did not fluidize the network, indicating that mere additional PLL binding is insufficient to drive the phase transition (Fig. 1J). These results confirm that the pre-formed coacervate droplet phase, which enables the coupled uptake of both ATP and PLL, is essential for mediating the solid-to-liquid transition of the condensates.

### Coacervate droplet infiltration mediates compositional diffusion and remodeling of RNA-peptide networks

Having established that ATP-PLL coacervates induce a solid-to-liquid phase transition of RNA aggregates, we sought to elucidate the underlying mechanism. We began by monitoring the phase transition process in real-time via confocal microscopy. We titrated fluorescently labelled ATP-PLL coacervates into a solution of pre-formed, Cy5-labelled rPU22-PLL aggregates (Fig. 2A, Supplementary Video 1). The transformation proceeded in two distinct stages: first, a macroscopic “shrinkage” of the solid aggregates, followed by a solid-to-liquid phase transition. This “transition” stage was characterized by a progressive increase in the local fluorescence intensity of RNA, ATP, and PLL within the condensates (Fig. 2A). The transformed condensates remain in a stable liquid state over extended timescales, resisting the aging and solidification typical of the original aggregates (Supplementary Fig. 6).

**Figure 2.**
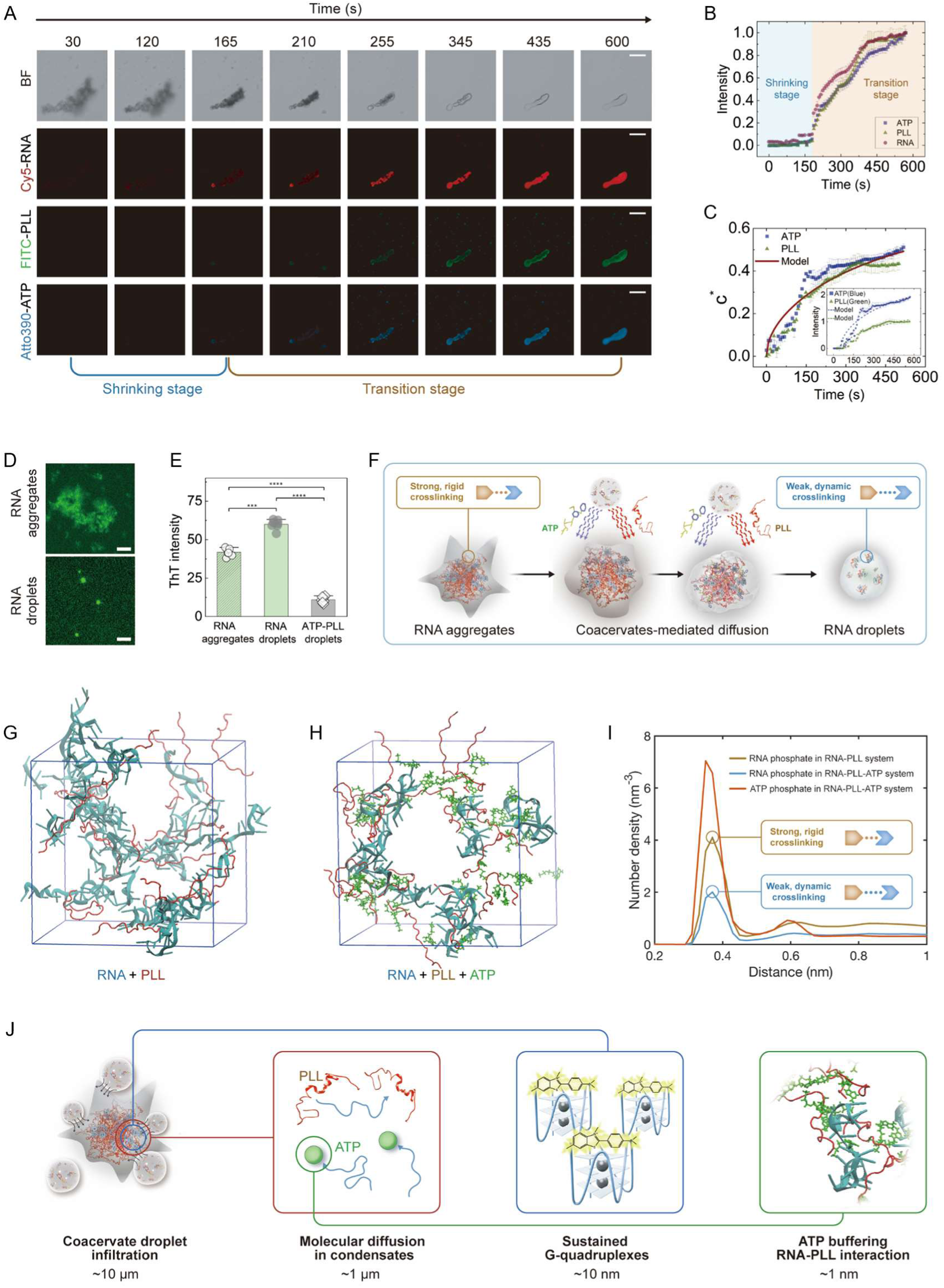
| The dynamics and molecular mechanisms of the coacervate-mediated solid-to-liquid phase transition. (**A**) Time-lapse confocal micrographs showing the co-diffusion of ATP (ATTO-390 labelled) and PLL (FITC labelled) from coacervates into solid RNA aggregates (Cy5-RNA added) upon contact. Scale bar, 50 µm. (**B**) Temporal evolution of fluorescence intensities for all three components during the transition. (**C**) Normalized concentrations of ATP and PLL within condensates during phase transition. Solid lines represent fits from a diffusion model, with normalized fluorescence intensities and model predictions shown in the inset. (**D**) Micrographs of Thioflavin T (ThT, 0.5 µM) assay within the initial aggregates and transformed RNA droplets. Scale bar, 10 µm. (**E**) Intensity of ThT fluorescence in RNA aggregates, RNA droplets as well as in ATP-PLL droplets. A two-tailed t-test was applied for statistical analysis with obtained P < 0.0001 (∗∗∗∗) and P < 0.001 (∗∗∗). (**F**) Schematic model for the co-diffusion of ATP and PLL buffering the static RNA-PLL interaction network, which enables fluidization but without disrupting RNA secondary structures. (**G**) All-atom molecular dynamic simulations conducted for RNA-PLL complexing system. (**H**) All-atom molecular dynamic simulations conducted for RNA-PLL-ATP ternary system. (**I**) Distribution of phosphate atoms (from RNA and ATP) around the ammonium groups of the PLL lysine residues. (**J**) Interpreting the multiscale mechanism of aggregate liquefaction via coacervates.

Quantitative analysis of fluorescence intensities corroborated this two-stage process. The initial shrinkage stage showed constant fluorescence levels, while the subsequent transition stage was characterized by a rapid, synchronous increase in all signals (Fig. 2B and Supplementary Fig. 7, measured intensity data). Since no external RNA was introduced into aggregates, the rising RNA signal indicates significant condensate compaction. To decouple molecular influx from volume changes, we normalized the ATP and PLL signals to the RNA signal (Fig. 2C, inset). The resulting synchronized influx curves for ATP and PLL suggest a co-diffusion mechanism. We therefore developed a diffusion model that accurately captures the kinetics of this co-influx, with the dimensionless concentration of ATP and PLL, *c*^∗^, expressed as

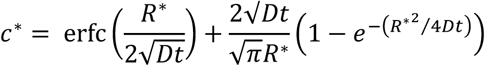

where *t* is time, *D* is molecular diffusion coefficients within condensates, *R*^∗^ represents the equivalent radius of aggregates, and erfc(*x*) denotes complementary error function (Supplementary text). The consistency of the experimental results with the diffusion model (Fig. 2C) indicates that the solid-to-liquid phase transition is driven by the co-diffusion of ATP and PLL from coacervate droplets into the solid aggregates.

Since rG4 structures promote aggregate solidification by enhancing RNA-PLL interactions, we asked whether the co-diffusion of ATP and PLL during the solid-to-liquid transition affects these rG4 structures. We probed rG4 integrity using thioflavin T (ThT), a G4-binding dye. The persistence of the ThT signal throughout the phase transition indicates that rG4 structures remain intact in both solid aggregates and the resultant liquid condensates (Fig. 2D, E). This demonstrates that the solid-to-liquid transition does not require the disruption of the RNA’s secondary structure.Taken together, we propose that the infiltrating ATP and PLL act as a molecular buffer, effectively weakening the multivalent, static cross-links between PLL and G-quadruplex RNA. This buffering action remodels the interaction network, transforming strong, rigid bonds into weaker, more dynamic interactions compatible with liquid-liquid phase separation (LLPS), as shown in Fig. 2F. Collectively, our data support a mechanism wherein the ATP-PLL coacervate phase serves as a delivery vehicle for the coupled influx of ATP and PLL. This co-diffusion enables a physical buffering of intermolecular interactions within the solid aggregate, leading to its fluidization without altering the intrinsic secondary structure of the RNA.

### ATP buffers RNA-peptide crosslinking to fluidize aggregates

The above results demonstrate that coacervate droplet-mediated delivery of ATP and PLL fluidizes RNA aggregates. To understand the underlying molecular mechanism, we performed all-atom molecular dynamics simulations of two systems: one containing RNA and PLL (ACE-K22-NME), and the other containing RNA, PLL (ACE-K22-NME), and ATP. Consistent with the macroscopic observations, the simulations revealed a direct competition mechanism. In the absence of ATP, RNA and PLL formed a system-spanning network via strong electrostatic interactions (Fig. 2G). The introduction of ATP disrupted this network; ATP molecules competed with RNA for binding sites on PLL, thereby inhibiting the formation of large-scale aggregates (Fig. 2H). This molecular picture supports the mechanistic model in Fig. 2F, wherein ATP diffusion into RNA aggregates converts strong, static RNA-PLL crosslinks into weak, dynamic interactions. To quantify this competition, we analyzed the radial distribution of phosphate atoms (from RNA and ATP) around the ammonium groups of PLL lysine residues. The distribution profiles (Fig. 2I) show that ATP involvement significantly reduces the local density of RNA phosphates around the PLL ammonium groups. Together, these simulation results indicate that ATP competitively binds to PLL, weakening the effective RNA-PLL interaction and fluidizing the aggregates. This molecular-scale evidence is consistent with the coacervate droplet-mediated diffusion process proposed from our experiments, establishing a framework for the multiscale mechanism of aggregate liquefaction via coacervates (Fig. 2J).

### Universal principle of coacervate-mediated solid-to-liquid phase transition

The simulations suggest that ATP introduced via coacervates acts as competitive binders by interacting with peptide to disrupt the static, multivalent RNA-peptide network, reshaping its interaction landscape from a kinetically trapped solid minimum into a thermodynamic equilibrium liquid basin (Fig. 3A). We therefore sought to determine if the resulting liquid condensates represent a thermodynamic equilibrium state or a kinetically trapped one. A key test was to examine the pathway dependence of the transition by varying the order of component mixing. Strikingly, the final liquid state was independent of the assembly pathway. Identical liquid droplets formed not only when pre-formed coacervates were added to pre-formed aggregates, but also when: (i) ATP and PLL were added separately to RNA aggregates (Fig. 3B), or (ii) all three components were individually mixed (Fig. 3C). FRAP analysis confirmed the fluidity of droplets generated via all pathways (Fig. 3B, C). This pathway independence demonstrates that the solid-to-liquid transition is a thermodynamically driven remodeling process, not a kinetic artifact.

**Figure 3.**
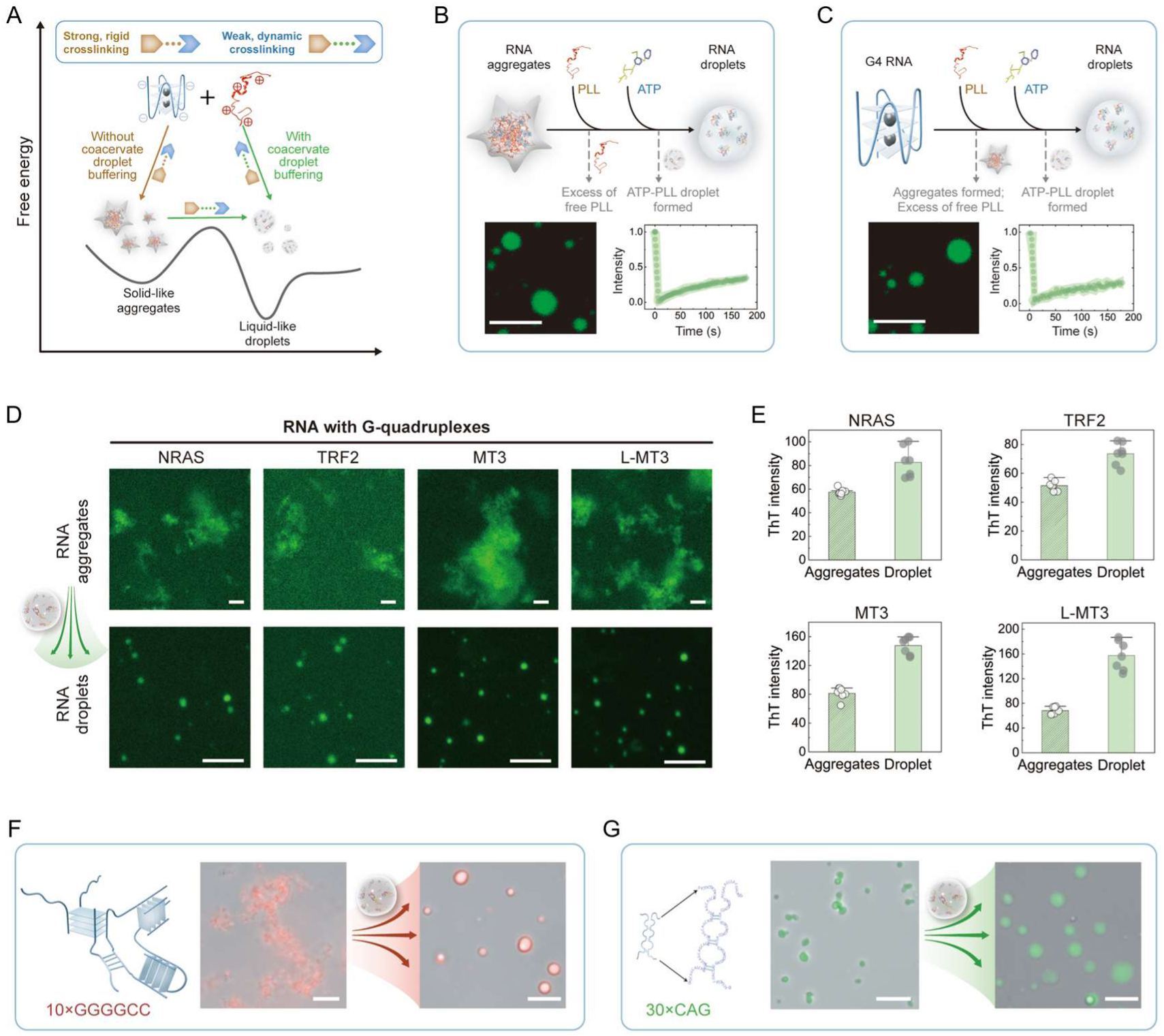
| Universal dissolution of RNA aggregates into liquid condensates via mixing with the ATP-PLL droplets. (**A**) Free energy landscape illustrating the coacervation and phase conversion of RNA-peptide complexes. The final liquid state is independent of the assembly pathway, converging on identical dissipative structures. (**B, C**) Identical liquid droplets form via two distinct pathways: (**B**) sequential addition of PLL then ATP to pre-formed solid aggregates, and (**C**) direct ternary co-assembly of all components. FAM-labeled rPU22 (green) visualizes distribution; Fluorescence Recovery After Photobleaching (FRAP) confirms liquid dynamics. All conditions maintain identical final stoichiometric concentrations. Scale bars: 10 µm. (**D**) Universal phase transition across solid-like aggregates formed from various G-quadruplex-forming RNAs (NRAS, TRF2, MT3, L-MT3) upon exposure to ATP-PLL coacervates. FAM-labeled RNAs were used for imaging. Scale bar: 10 µm. (**E**) Corresponding ThT fluorescence assays confirm the structural preservation of G-quadruplex motifs throughout the phase transition for all RNA types shown in (**D**). (**F, G**) Functional relevance demonstrated by the dissolution of pathologically relevant solid-like aggregates formed by repeat expansion RNAs: (**F**) [GGGGCC]₁₀ (Cy5-labeled) and (**G**) [CAG]₃₀ (FAM-labeled). Both undergo a solid-to-liquid transition upon ATP-PLL coacervate addition. Scale bar: 10 µm.

To define the conditions enabling this transition, we systematically varied the stoichiometry of ATP-PLL droplets and RNA aggregates. The resulting phase diagram (Supplementary Fig. 8) shows the transition is robust, occurring across a broad range of stoichiometries. However, when the amount of RNA aggregates is comparable to or exceeds that of the ATP-PLL droplets, the phase transition is inhibited, leaving residual solid aggregates. This stoichiometric threshold is consistent with the competitive binding mechanism; a sufficient concentration of ATP relative to RNA is required to shift the interaction mode and drive the system to a homogeneous liquid equilibrium state.

With the established mechanism by which ATP-PLL coacervates fluidize RNA aggregates, we next investigated the generality of this coacervate-mediated solid-to-liquid transition. We hypothesized that the process, driven by the buffering of molecular interactions rather than disrupting specific RNA folding structures, should be applicable to a broad class of RNA-peptide solids. We first tested this on a panel of diverse RNA sequences known to form stable G-quadruplex structures (NARS, TRF2, MT3, as well as its chiral counterpart L-MT3). All formed solid-like aggregates with PLL (Fig. 3D). Strikingly, upon addition of ATP-PLL coacervates, every one of these RNA-PLL aggregates underwent a complete transition, forming spherical droplets (Fig. 3D). Critically, ThT fluorescence confirmed that the G4 structures remained intact in both the initial solids and the resultant liquid droplets (Fig. 3E). This universal preservation of RNA secondary structure across different sequences strongly reinforces our model that the transition is mediated by a physical buffering of intermolecular crosslinking, not by the disruption of RNA folding. We further extended our analysis to pathologically relevant RNAs with repeat expansions, including 10×GGGGCC (Fig. 3F) and 30×CAG (Fig. 3G). These RNAs, which also form solid-like aggregates with PLL, similarly underwent a complete solid-to-liquid transition upon exposure to ATP-PLL coacervates. The consistent fluidization of aggregates formed by structurally diverse RNAs, from short, structured G4 motifs to longer, repetitive sequences, demonstrates that the coacervate-mediated transition is a universal phenomenon, largely independent of RNA length and specific sequence.

### Composition-dependent control of liquefaction efficacy

We next asked whether the observed phase transition is a unique property of ATP-PLL coacervates or a more general phenomenon. To this end, we systematically tested a library of coacervates with diverse compositions. The efficacy in inducing the solid-to-liquid transition of rPU22-PLL aggregates revealed a clear hierarchy, allowing us to categorize the coacervates into three distinct classes based on aggregate miscibility (Fig. 4A). Specifically, coacervates such as ADP (adenosine diphosphate)-PLL and ATP-PLR (Poly-L-arginine) generated homogeneous, liquid droplets upon mixing with RNA aggregates, indicating ‘fully miscible’ scenario of the solid aggregate, analogous to ATP-PLL coacervates. Coacervate systems containing long-chain RNA homopolymer, like rA_100_-PLL and Spermidine-rA_100_ induced the formation of spherical condensates, but these exhibited inhomogeneous fluorescence inside, suggesting ‘partially miscible’ situations with limited component exchange. The surface plot of the fluorescence intensity within the condensate (Fig. 4B) further supports the distinction between the “Fully miscible” and “Partially miscible” categories: the former exhibits a paraboloid distribution, whereas the latter shows an irregular, multi-peaked profile. In stark contrast, coacervates formed by complexing of rA_100_ and rU_100_ with K_72_, a supercharged protein^36^, and by complexing of PAA (Polyacrylic acid) with PDDA (Poly(diallyldimethylammonium chloride)), showed no interaction, leaving the solid-like aggregates unchanged and corresponding to the ‘immiscible’ condition. Notably, coacervates in the immiscible group are characterized by intrinsically weak electrostatic interactions. This is evidenced by their lower salt resistance compared to other groups (Table. S1). In particular, the low charge density of K_72_ prevents its coacervation with rA_100_ or rU_100_ alone. In this system, phase separation only occurs in the ternary mixture (rU_100_-rA_100_-K_72_), where A-U base pairing enables the two RNA strands to hybridize, forming a combined polyanion of effectively higher charge density that can interact with K_72_.

**Figure 4.**
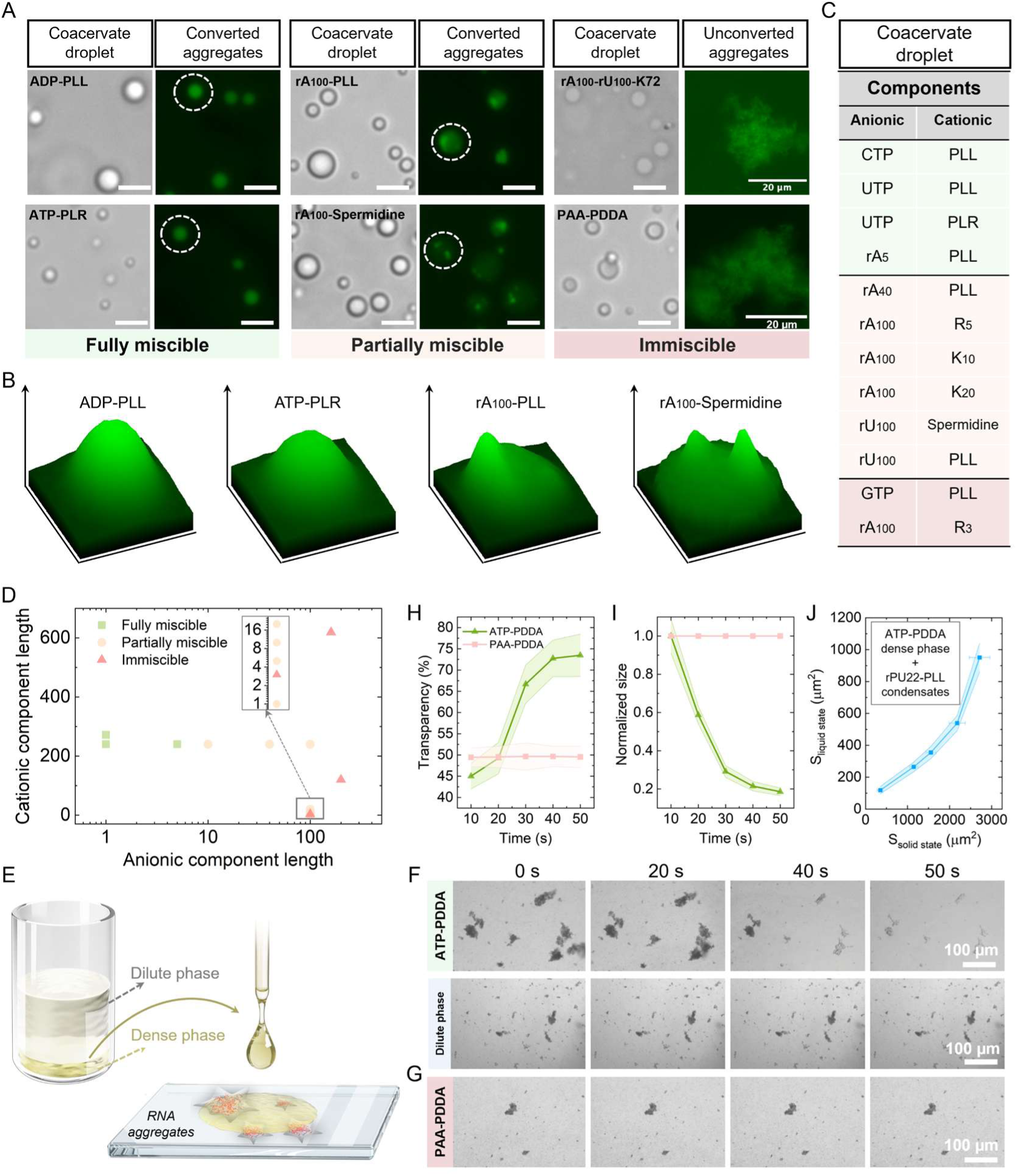
| Selective solid-to-liquid phase transition of RNA-PLL aggregates via molecularly tunable coacervate compartments. **(A)** Coacervate systems exhibit composition-dependent miscibility with solid RNA-PLL aggregates, classified as fully miscible, partially miscible, or immiscible. Brightfield images show coacervate droplets; fluorescence images show FAM-labeled rPU22 distribution after introducing droplets into aggregates. Scale bar: 5 μm (unless noted). **(B)** Surface plots of fluorescence intensity within condensate regions marked in (A), highlighting RNA spatial distribution features in “fully miscible” and “partially miscible” groups. (C) Screening of coacervate compositions confirms the miscibility categories in (A). (**D**) The influence of cationic and anionic polymer length on aggregate liquefaction efficiency. (**E)** Macroscopic dissolution assay demonstrating phase transition of RNA aggregates via bulk dense phase of coacervates. **(F)** ATP-PDDA coacervate dense phase dissolves solid RNA-PLL aggregates while dilute phase has no dissolution capacity. **(G)** PAA-PDDA dense phase shows no dissolution capacity. **(H, I)** Transparency and size changes of RNA aggregates upon addition of ATP-PDDA versus PAA-PDDA dense phases. **(J)** Area of transformed, liquid-state RNA-PLL condensates as a function of initial solid aggregate area, following exposure to ATP-PDDA coacervates. Abbreviations: ADP, adenosine diphosphate; AMP, adenosine monophosphate; CTP, Cytidine triphosphate; UTP, uridine triphosphate; GTP, Guanosine triphosphate; PLL, Poly-L-lysine; PLR, Poly-L-arginine; rA_5_, 5-mer adenosine homoribonucleotide; rA_40_, Polyadenylic acid (adenine homopolymer, 40 nt); rA_100_, Polyadenylic acid (100 nt); rU_100_, Polyuridylic acid (uracil homopolymer, 100 nt); K_72_, Supercharged cationic protein containing 72 lysine residues; PAA, Polyacrylic acid; PDDA, Poly(diallyldimethylammonium chloride); R_3_, Tripeptide (Arg-Arg-Arg); R_5_, Pentapeptide (Arg-Arg-Arg-Arg-Arg); K_10_, Lysine decapeptide; K_20_, 20-mer lysine oligopeptide.

Our results indicate that the efficacy in inducing solid-to-liquid phase transitions is closely correlated with the molecular characteristics of the coacervate’s component. The most effective (“Fully miscible”) coacervates were consistently formed by highly charged polycations paired with small-molecule anions like ATP or ADP. This observation aligns precisely with a diffusion-dominated buffering mechanism, wherein coacervate droplets efficiently infiltrate aggregates and the high mobility of the small anions enables them to compete with RNA for polycation (e.g. PLL and PLR within aggregates) binding. This potent buffering action effectively disrupts the static rPU22-PLL network, facilitating its fluidization. Coacervates classified as “Partially miscible” (e.g., rA_100_-PLL coacervates) possess sufficient cohesive interaction strength to infiltrate the aggregate interface. However, their capacity for network remodeling is limited by the restricted molecular mobility of the long-chain anions, whose slow diffusion kinetics impede rapid competition with the pre-existing, multivalent RNA-peptide crosslinks. This results in an incomplete fluidization. Conversely, systems in the “Immiscible” regime fail at the initial infiltration step. Coacervates formed from polyelectrolyte pairs with intrinsically weak interaction enthalpies (e.g., rU_100_-rA_100_-K_72_ or PAA-PDDA) lack the cohesive driving force necessary to partition into and wet the dense aggregate phase. This is analogous to individual ATP or PLL molecules, which cannot enter aggregates alone (Fig. 1I, J). Thus, a sufficiently high interaction strength among coacervate constituents is a prerequisite to mediate their collective infiltration as a condensed phase. Without this infiltration, no dynamic exchange is possible, leaving the static network intact.

This molecular framework provides a mechanistic basis for the observed hierarchy of coacervate efficacy, governed by two independent design parameters: interaction strength for aggregate infiltration and constituent mobility for internal rearrangement. To translate this mechanism into a predictive design rule, we screened a panel of complex coacervates (Fig. 4C). The data robustly validate a two-parameter model where infiltration requires sufficient coacervate cohesion, and liquefaction requires high anionic mobility. Only coacervates formed by small, mobile anions with highly charged polycations, such as UTP (uridine triphosphate)-PLR and rA_5_-PLL coacervates, meet both criteria, achieving complete liquefaction (Fig. 4C, Supplementary Fig. 9A). Systems using longer anionic polymers (rA_40_, rA_100_, rU_100_) satisfy the infiltration requirement but fail the mobility criterion, resulting in partial miscibility (Fig. 4C, Supplementary Fig. 9B). This delineation explicitly identifies anionic constituent within the coacervate as the critical parameter controlling liquefaction efficiency (Fig. 4D). The framework also explains edge-case failures. The rA_100_-R_3_ (Arg-Arg-Arg tripeptide) pair lacks sufficient interaction strength to form a stable condensate, failing the infiltration prerequisite. Interestingly, GTP-PLL mixtures form solid-like condensates themselves due to auxiliary Hoogsteen base-pairing; this intrinsic rigidity rendering them ineffective as liquefying agents (Fig. 4C, Supplementary Fig. 9C).

### Bulk dissolution of aggregates via coacervate liquefiers

The general applicability of using coacervates composed of small molecules to drive the solid-to-liquid transition of RNA aggregates indicates that these coacervates can serve as “effective solvent” to liquefy RNA aggregates. To test this hypothesis on a macroscopic scale, we performed a bulk dissolution assay (Fig. 4E). When solid rPU22-PLL aggregates were placed in contact with the dense coacervate phase of ATP-PDDA, they shrank and dissolved over time (Fig. 4F, Supplementary Video 1). In stark contrast, aggregates exposed to the dilute phase of ATP-PLL coacervate mixture, or a PAA-PDDA dense phase remained stable (Fig. 4F, G, and Supplementary Video 3). Quantitative analysis confirmed that the ATP-PDDA phase increased the optical transparency (Fig. 4H) and decreased the aggregate size (Fig. 4I, J), whereas the PAA-PDDA phase produced no significant change (Fig. 4H, I). This macroscopic demonstration further confirms that the small-molecule anionic species, such as ATP, is the decisive factor enabling coacervates to drive the solid-to-liquid phase transition of RNA aggregates.

Altogether, we establish a mechanistic framework in which coacervate droplets mediate the diffusion of their constituent molecules into RNA-peptide aggregates. This diffusion is suppressed in the absence of coacervates, for example, when only the constituent molecules are present. With this coacervate-mediated diffusion process, the key determinant to trigger the solid-to-liquid phase transition is the presence of small-molecule anionic component, which functions as a dynamic molecular buffer to remodel the RNA-peptide interaction network inside the aggregates.

### Functional rescue of RNA by coacervate-mediated liquefaction

Our finding that coacervates can buffer and remodel the internal interaction network of solid-like aggregates, together with the correlation between a condensate’s internal interaction network and its function as a biomolecular compartment^29^, led us to probe if such coacervate-mediated liquefaction mechanism could be harnessed to control RNA structure and function. To test this, we first employed an RNA molecular beacon^37^, which fluoresces only when its hairpin structure is disrupted (Fig. 5A). We found that sequestration in solid rPU22-PLL aggregates locked it in a static and folded “OFF” state. Strikingly, coacervate-mediated liquefaction triggered strong fluorescence within the resulting liquid droplets, with intensity increasing ∼5-fold (Fig. 5B, C). This demonstrates that the transition remodels the RNA’s environment from a constrained solid to a dynamic liquid, enabling structural reconfiguration.

**Figure 5.**
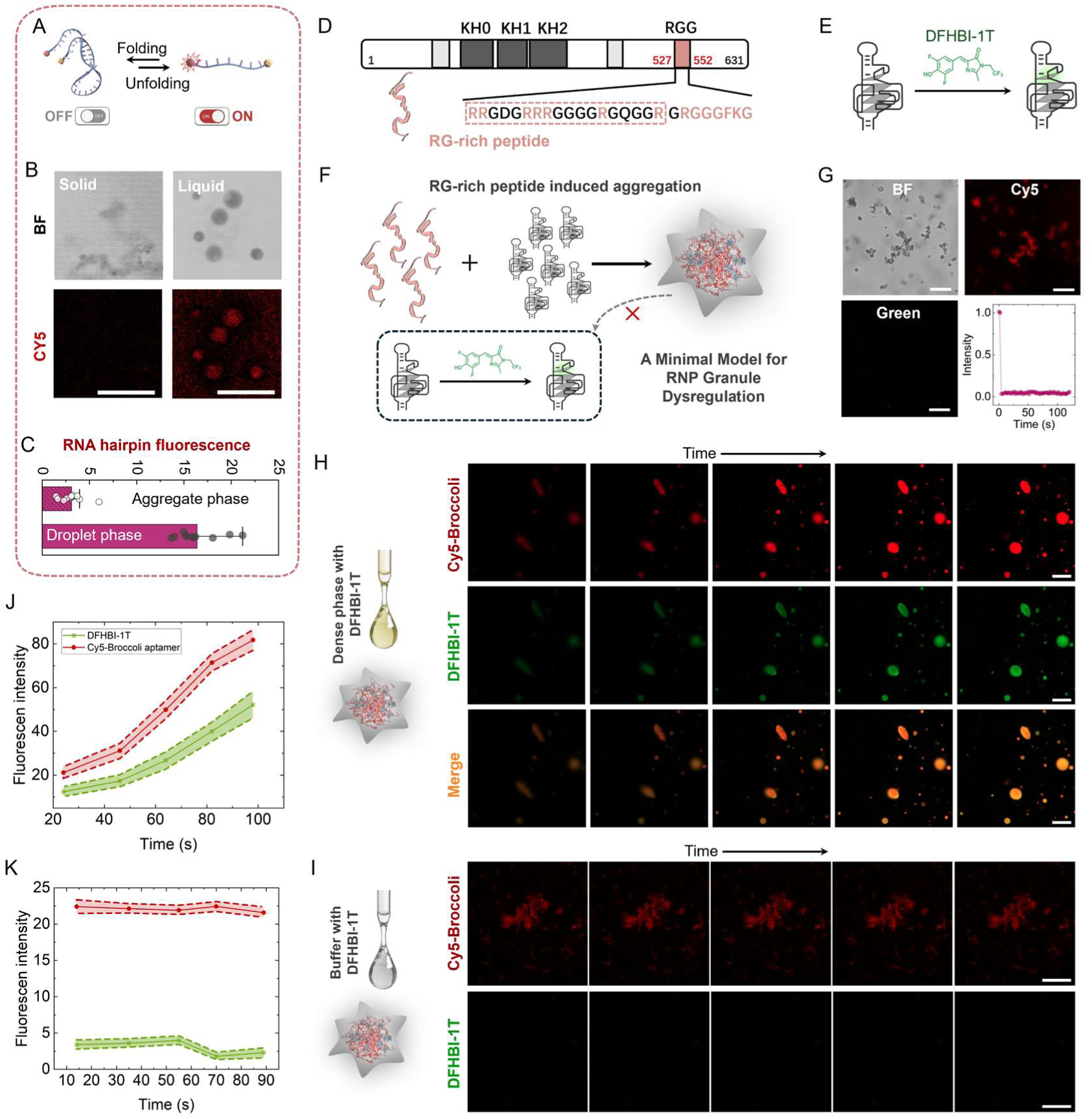
| Coacervate-mediated liquefaction rescues RNA function from aggregation-induced quenching. (**A**) Schematic of the RNA molecular beacon assay. Fluorescence is quenched in the native hairpin state and emitted upon unfolding. (**B**) Confocal micrographs show the molecular beacon is quenched within solid rPU22-PLL aggregates but fluoresces after coacervate-induced transition to liquid droplets. Scale bar: 10 µm. (**C**) Quantification of fluorescence intensity in the solid aggregate versus liquid droplet phases. (**D**) Domain schematic of the arginine-glycine-rich (RGG) peptide derived from Fragile X Mental Retardation Protein (FMRP). (**E**) Schematic of the Broccoli RNA aptamer assay. Fluorescence requires correct RNA folding to bind and activate the small-molecule fluorophore DFHBI-1T. (**F**) Cationic RGG peptide drives Broccoli aptamer into solid-like aggregates as a minimal model for ribonucleoprotein (RNP) granule dysregulation. (**G**) Cy5-labeled Broccoli aptamer aggregates formed with the RGG peptide. Scale bar: 20 µm. (**H**) Functional rescue by coacervate liquefier. Addition of ATP-PDDA coacervates (dense phase) containing DFHBI-1T triggers solid-to-liquid transition and concurrent emergence of DFHBI-1T fluorescence, indicating aptamer refolding and fluorophore activation. Scale bar: 20 µm. (**I**) Control experiment: addition of DFHBI-1T in buffer (without coacervates) fails to induce phase transition and fluorescence recovery. Scale bar: 50 µm. (**J, K**) Quantitative analysis of the processes in (**H**) and (**I**). Cy5 fluorescence intensity (reporting Broccoli aptamer localization) and DFHBI-1T fluorescence intensity (reporting functional aptamer-fluorophore binding) show that functional recovery is tightly coupled to coacervate-mediated liquefaction.

To further demonstrate the functional utility of coacervate-mediated liquefaction, we posed a more targeted utilization with practical implications: using the coacervate liquefier to rescue functional RNA from a sequestered, quenched state. To this end, we selected a cationic, arginine-glycine-rich (RGG-rich) peptide motif derived from Fragile X Mental Retardation Protein (FMRP, Fig. 5D), and the Broccoli RNA aptamer, which binds and fluorescently activates the small-molecule fluorophore DFHBI-1T upon proper folding (Fig. 5E)^38^. Functional silencing and quenching of RNA via cationic protein-induced aggregation (Fig. 5F) is a widespread phenomenon, relevant to RNA dysregulation in diseases^39^ and synthetic cells^40^. To directly visualize this quenching process, we employed a Broccoli aptamer labeled with Cy5, enabling independent tracking of RNA localization and functional activation (via green fluorescence of DFHBI-1T). Mixing the Cy5-Broccoli RNA with the RGG-rich peptide drove the formation of solid-like aggregates, which sequestered the RNA and rendered its mobility (Fig. 5G).

Upon addition of a dense phase of ATP-PDDA coacervates pre-loaded with DFHBI-1T, the aggregates underwent a solid-to-liquid transition, with increasing of Cy5-Broccoli aptamer fluorescence akin to those in Fig. 2A. Concurrently, strong green fluorescence emerged from the resulting phase transition, indicating successful infiltration of DFHBI-1T and its activation by the now-accessible RNA aptamer (Fig. 5H). In a critical control, adding DFHBI-1T alone (without coacervates) produced neither phase transition nor fluorescence recovery, confirming that the solid aggregate matrix itself kinetically traps and functionally silences the encapsulated RNA aptamer (Fig. 5I). Quantitative analysis solidified these observations. The fluorescence intensity of both Cy5-Broccoli aptamers and Broccoli-fluorophore complexes increased in direct correlation with the degree of liquefaction (Fig. 5J), while control aggregates showed no fluorescence changes, indicating the absence of RNA function activation (Fig. 5K).

Together, these experiments establish a principle that the coacervate liquefier can be deployed to reverse functional quenching of RNA within aggregates. By competitively buffering peptide-RNA interactions, the liquefier dismantles diffusion barriers, liberates RNA, and restores the dynamic milieu necessary for folding and function (Fig. 5H). Fundamentally, this suggests a potential cellular strategy, employing endogenous “liquefier” condensates to regulate RNP granules. Technologically, it provides a versatile tool for reactivating aggregated biologics, rescuing RNA functionality in synthetic cells, where controlling phase behavior is synonymous with controlling function.

## Discussion

In this work, we have discovered a fundamental principle by which liquid-like coacervates can trigger the solid-to-liquid phase transition of a wide range of kinetically trapped RNA-peptide aggregates. This process is driven by the coacervate’s capacity to act as a source of mobile molecular buffers, that infiltrate and reorganize the static network of the solid aggregate. Once inside, these molecules competitively buffer the original multivalent interactions between RNA and peptides, reshaping the free energy landscape and collectively driving the system into a dynamic, liquid state.

Our findings establish several key design principles for the liquefaction of solid-like aggregates. First, the mechanism is general, applicable to a broad range of RNA aggregates stabilized by RNA intra– or intermolecular hybridization. Second, the efficacy of the “catalytic” coacervate is not generic but is exquisitely dependent on the molecular nature of its components, with small, mobile anions such as ATP are uniquely effective. Third, this phase transition is not merely a material change but is functionally coupled to the folding state of encapsulated RNA, with the resultant liquid condensates serving as a switch to unfold the RNA hairpin and rescue Broccoli aptamer functionality.

Conceptually, this mechanism draws an analogy to the action of molecular chaperone and RNA buffering effects for the inhibition of protein aggregates^17, 41^. Here, the coacervate itself functions as a chaperone, to reverse aberrant RNA aggregates into dynamic liquid states. This establishes a new paradigm for complex coacervates, moving beyond their role as passive compartments to active media for the liquefaction of solid biomolecular assemblies. Consequently, this work suggests that a sophisticated hierarchy may exist within cells, where one condensate can regulate the material state of another, particularly for those multiphase biomolecular condensates^42^. By providing a physical mechanism for such interactions, our study not only advances the fundamental understanding of biomolecular phase behavior but also opens novel therapeutic avenues. The targeted dissolution of pathological aggregates, a hallmark of neurodegenerative disease, through engineered condensate catalysts represents a promising frontier inspired by these principles^5^.

## Materials and Methods

Detailed descriptions of materials, experimental procedures, theoretical analysis, and molecular dynamics simulations are provided in the Supplementary Information.

## Data availability

The data that support this study are available within the paper and its Supplementary Information. Further details and raw images are also available from the corresponding authors upon reasonable request.

## Code availability

The input files and analysis scripts for all-atom molecular dynamics simulations are available from the corresponding authors upon reasonable request.

## Acknowledgments

We thank Mr. Kwan Kiu Lau for his contributions during the initial phase of this project and Dr. Gurudas Chakraborty and Mr. Fabian Wiertz from Prof. Andreas Herrmann’s team for providing the supercharged protein sample. We are grateful to Prof. Howard Stone and Prof. Tiantian Kong for insightful discussions. We acknowledge support by the General Research Fund (Nos. 17306221, 17317322) from the Research Grant Council (RGC) of Hong Kong, RGC Senior Research Fellow Scheme (SRFS2425-7S04). H.C.S. was funded in part by the Croucher Senior Research Fellowship from Croucher Foundation and the Health@InnoHK program of the Innovation and Technology Commission of the Hong Kong SAR Government. W.G. acknowledge funding support by Shenzhen University (000008021123, 000008010352). All-atom molecular dynamics simulations were performed on MGPU cluster at the High Performance Computing Center (HPCCC) of Hong Kong Baptist University.

## Author contributions

W.G. conceived and designed the study. W.G. and R.L. conducted the experiments. X.Z. performed the molecular simulations. W.G., X.Z. and Z.L. analyzed and interpreted the data. W.G. wrote the original draft. Y.S., X.Z., Z.L., and H.C.S. provided critical feedback and contributed to manuscript revision. H.C.S. acquired funding and supervised the project. All authors reviewed, edited, and approved the final manuscript.

